# *Leishmania major* targets macrophage Syntaxin-2 to impair phagolysosome biogenesis and promote intracellular survival

**DOI:** 10.64898/2026.06.28.735094

**Authors:** Suman Samanta, Arghyajyoti Pramanik, Rupak Datta, Subhankar Dolai

## Abstract

Macrophages destroy pathogens by engulfing them into phagosomes that mature into degradative phagolysosomes via lysosome fusion. *Leishmania* parasites subvert this antimicrobial pathway to establish intracellular infection and cause leishmaniasis. We previously identified the SNARE protein syntaxin-2 (Stx2) as a promoter of phagolysosome biogenesis that simultaneously limits particle binding and uptake. Consistent with this dual role, Stx2-depleted macrophages (Stx2-KD) show enhanced binding and internalization of *Leishmania major*. Stx2-KD macrophages also sustain higher intracellular parasite loads. We find that *L. major* actively targets macrophage Stx2 by selectively depleting Stx2 from phagosomes through its virulence metalloprotease GP63. Phagosomes containing GP63-deficient *L. major* retain Stx2 and acquire increased levels of lysosomal hydrolases and v-ATPase, restoring degradative capacity. In BALB/c mice, *L. major* infection markedly reduces Stx2 in infected tissues in a GP63-dependent manner. Collectively, our findings identify GP63-mediated Stx2 depletion as a key virulence strategy of *L. major*, positioning the GP63-Stx2 axis as a promising therapeutic target for leishmaniasis.

## INTRODUCTION

Leishmaniasis, which ranges from cutaneous to fatal visceral disease, remains endemic in nearly 100 countries and lacks effective vaccines or safe therapies (Pareyn et al., 2025). Infection is initiated when *Leishmania* promastigotes transmitted by sand flies are internalized by macrophages into phagosomes (Podinovskaia and Descoteaux, 2015). Phagosomes normally mature through sequential fusion with endolysosomal compartments to generate acidic, degradative phagolysosomes that kill pathogens and process antigens (Flannagan et al., 2012). *Leishmania* promastigotes instead arrest phagolysosome biogenesis, enabling their differentiation into phagolysosome-adapted amastigotes that thrive and proliferate within phagolysosome-like parasitophorous vacuoles (PVs) (Chang and Dwyer, 1976; Moradin and Descoteaux, 2012; Young and Kima, 2019). Identifying host machinery targeted by *Leishmania* to block phagolysosome biogenesis is therefore key to understanding parasite survival and restoring macrophage microbicidal function.

SNARE proteins drive fusions for acquiring endolysosomal compartments during phagosome biogenesis and maturation (Desjardins et al., 1994; Flannagan et al., 2012). Q-SNAREs (syntaxins/SNAPs) and R-SNAREs (VAMPs or Sec22) on opposing membranes assemble into trans-SNARE complexes, with syntaxin serving as central organizers (Jahn and Scheller, 2006; Sudhof and Rothman, 2009). *Leishmania* PVs acquire non-lytic ER and ER–Golgi compartments via the ER/Golgi SNAREs syntaxin-5, syntaxin-18, and Sec22b (Canton and Kima, 2012; Canton et al., 2012). Further, parasites exclude phagosomal syntaxin-activator synaptotagmin v by lipophosphoglycan (Vinet et al., 2011), and use GP63 to cleave VAMP8, impairing NADPH oxidase-dependent phagosome alkalization and antigen processing (Delamarre et al., 2005; Matheoud et al., 2013). However, whether *Leishmania* directly targets phagosomal syntaxins that mediate lysosome acquisition remains absolutely unknown.

Among phagosomal syntaxins (Hackam et al., 1996), syntaxin-3 and syntaxin-4 promote cytokine secretion by recruiting recycling endosomes to nascent phagosomes (Stow et al., 2006; Verboogen et al., 2017). In contrast, syntaxin-2 (Stx2) drives phagosome maturation by enhancing phagosome–lysosome fusion while simultaneously restricting non-lytic compartment acquisition and surface recycling of phagocytic receptors (Samanta et al., 2025b). Stx2 thus represents a promising host target for intracellular pathogens, and its loss in macrophages likely facilitates both pathogen uptake and intracellular survival. Consistently, we found that Stx2 depletion (Stx2-KD) in macrophages increases uptake and survival of cutaneous leishmaniasis-causing pathogen *L. major*. In transient macrophage infection, *L. major* also depletes phagosomal Stx2 but more robust in infected tissues via GP63. Thus *Leishmania* exploits Stx2 depletion to subvert phagosome maturation and establish infection.

## RESULTS

### Stx2 depletion enhances *L. major* uptake and increased intracellular parasite burden in macrophages

Stx2-KD macrophages showed increased engagement and phagocytosis but impaired degradation of IgG-opsonized particles (Samanta et al., 2025c), a condition favorable for infection and survival of macrophage dwelling pathogens. To verify this we used cutaneous leishmaniasis causing pathogen *L. major* that infect macrophages via phagocytosis. We initiated infection by incubating *L. major* promastigotes with macrophages. Scanning electron microscopy revealed increased association of *L. major* promastigotes with Stx2-KD macrophages within 10 min of infection, indicating enhanced early parasite binding (Fig. 1 A,B). To examine internalization of parasites, macrophages incubated with *L. major* for 12 hr were subjected to inside-outside staining by using anti-GP63 antibody to selectively label externally exposed *L. major*. Confocal analysis revealed significantly increased numbers of internalized parasites in 12 h post-infected Stx2-KD macrophages. These observations align with our previous work (Samanta et al., 2025c) showing that Stx2 limits surface recycling of Fc receptors for phagocytosis. Stx2-KD likely also increases surface availability of other phagocytic receptors for *Leishmania* uptake, thereby promoting parasite attachment and internalization.

**Figure 1.**
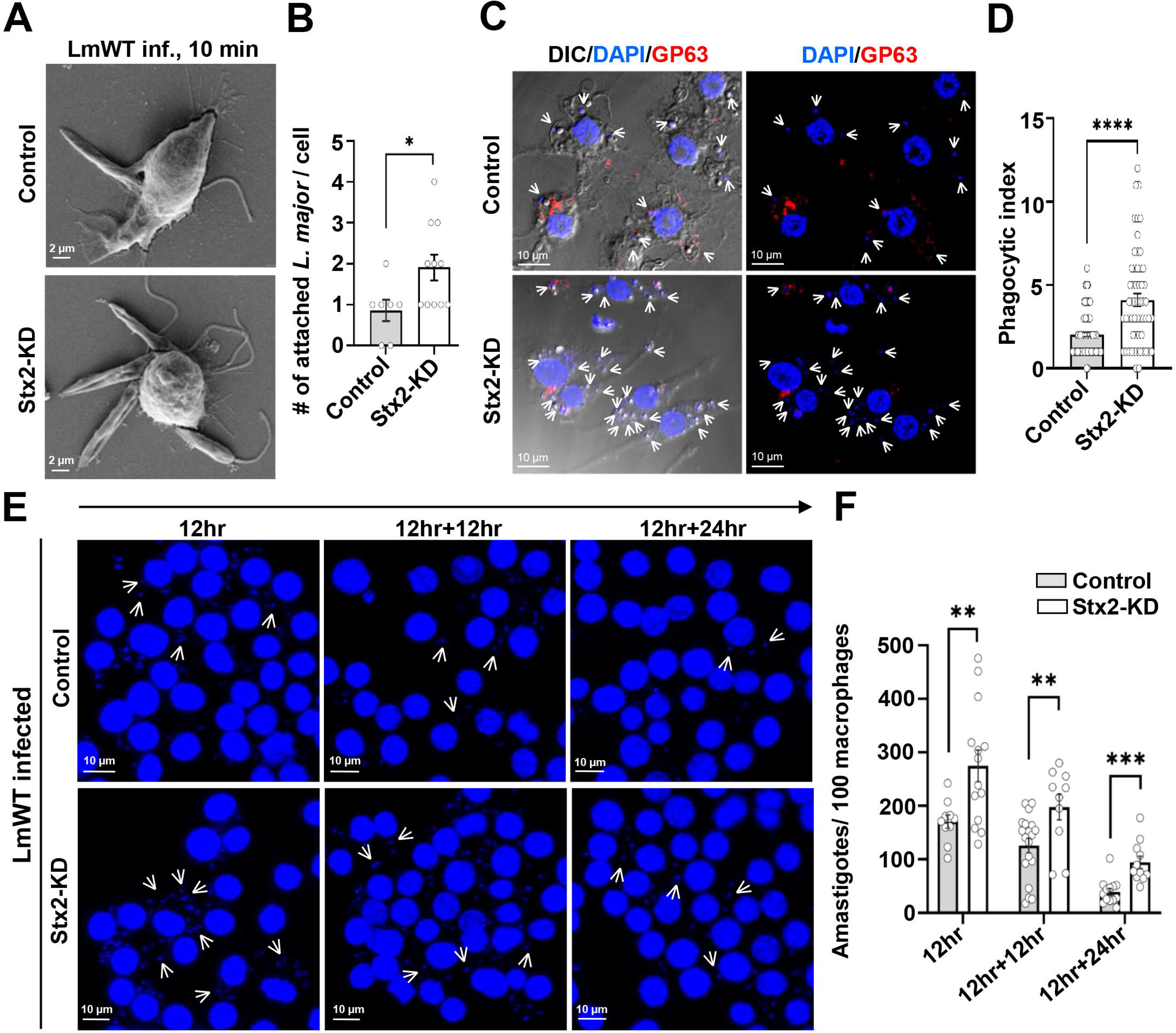
Stx2 depletion enhances *Leishmania major* uptake and promotes intracellular survival, increasing parasite burden in macrophages. (A) Representative scanning electron microscopy (SEM) images of control and Stx2-knockdown (Stx2-KD) macrophages infected with *L. major* for 10 min, showing parasite association at the cell surface. (B) Quantification of surface-attached parasites per macrophage (control, n = 7; Stx2-KD, n = 11). Data represent mean ± s.e.m. from three independent experiments; *P < 0.05. (C) Representative confocal maximum intensity projection images of macrophages infected with *L. major* for 12 h, showing differential inside (unstained) and outside (red) labeling using anti-GP63 antibody. Nuclei are stained with DAPI (blue). Parasite nuclei are indicated by white arrowheads. Scale bars: 10 μm. (D) Quantification of phagocytic index. Data represent mean ± s.e.m. from three independent experiments, with individual values shown; ****P < 0.0001. (E) Representative immunofluorescence images of intracellular parasite burden in control and Stx2-KD macrophages at 12 h post-infection. For extended time points (12 h + 12 h and 12 h + 24 h), extracellular parasites were removed after 12 h, followed by further incubation for 12 or 24 h. Nuclei are stained with DAPI (blue), and parasite nuclei are indicated by white arrows. (F) Quantification of parasite burden, expressed as the number of amastigotes per 100 macrophages. At least 100 macrophages were analyzed per condition. Data represent mean ± s.e.m. from three independent experiments; **P ≤ 0.01; ***P ≤ 0.001 (unpaired two-tailed Student’s t-test).

We next examined whether Stx2 depletion impacts intracellular parasite survival. Macrophages were infected with pathogens for 12 h for sufficient internalization, and after removing unbound and loosely bound parasites, further incubated for additional 12 and 24 h. Confocal detection of DAPI-stained *L. major* nuclei (pointed by white arrowheads) revealed marked increase in intracellular parasite burden at 12 h, which remained persistently higher upon extended incubation (12 h + 12 h and 12 h + 24 h) (Fig. 1 E,F). These data show increased parasite load in Stx2-KD macrophages is accompanied with improved intracellular survival (Fig. 1 E,F). Collectively, these results indicate that loss of Stx2 not only facilitates parasite entry but also supports intracellular survival.

### *Leishmania major* depletes phagosomal Stx2 via its virulence factor GP63

Evasion of lysosome-mediated destruction is a strategy adopted by intracellular pathogens (Liehl et al., 2015). For *Leishmania* parasites, cessation of phagolysosome biogenesis is critical for securing early promastigote survival in macrophages (Moradin and Descoteaux, 2012). Since, macrophage phagosome-enriched SNARE Stx2 drives lysosome acquisition during phagolysosome biogenesis (Hackam et al., 1996; Samanta et al., 2025c), we asked whether *L. major* modulates Stx2 during infection. RAW 264.7 macrophages were challenged with IgG-opsonized inert zymosan particles or infected with *L. major* promastigotes and localization and expression of Stx2 were examined by confocal immunofluorescence imaging (Fig. 2). Lysosomal LAMP1 was also probed to delineate phagosomal membranes (Czibener et al., 2006; Fig. 2A). In resting macrophages, Stx2 displayed consistent localization on the plasma membrane and intracellular vesicles (Fig. 2A, Fig. S1) as observed earlier (Samanta et al., 2025c). Stx2 was found to localize robustly around zymosan-containing phagosomes, and remained associated throughout the time course of study (Fig. 2A). In contrast, *L. major*-containing phagosomes showed markedly reduced Stx2 levels beginning at 6 h post-infection (p.i.) (Fig. 2A, indicated by arrowhead). Quantification of phagosomal Stx2 fluorescence intensity revealed maximal depletion at 6 h p.i., which persisted through 12 h p.i. before recovering by 24 h p.i. (Fig. 2B). Notably, immunoblotting revealed no change in total cellular Stx2 levels, and no cleavage products were detected (Fig. 2C,D). This indicates selective exclusion of Stx2 from parasite-containing phagosomes rather than proteolytic degradation (Fig. 2C,D). While pathogens are known to subvert phagosome maturation by substituting SNARE partners (Santamaria et al., 2025) or deploying SNARE-mimetic effectors (Singh et al., 2018), our findings provide the first evidence of pathogen-induced exclusion of a phagosomal SNARE. This represents a distinct mechanism by which *L. major* stalls phagolysosome biogenesis to promote intracellular survival.

**Figure 2.**
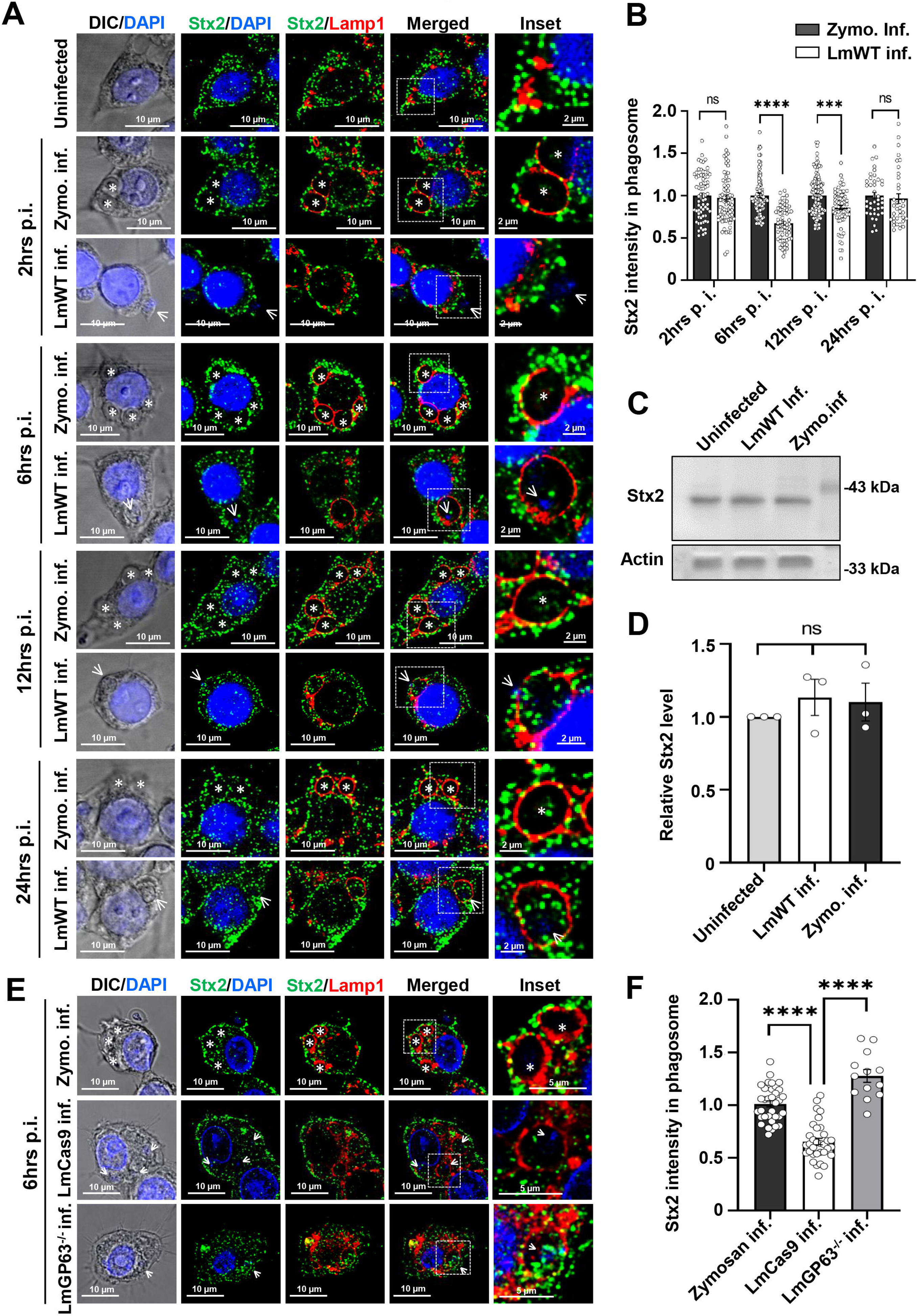
GP63-dependent depletion of phagosomal Stx2 by *Leishmania major* in macrophages. (A) Representative DIC and confocal immunofluorescence images of RAW264.7 macrophages stained for endogenous Stx2 (green) and Lamp1 (red) under control conditions or following uptake of opsonized zymosan particles or infection with *L. major* for 2, 6, 12, and 24 h. Nuclei were stained with DAPI (blue). Parasite nuclei are indicated by white arrows and zymosan particles by white asterisks. Insets show magnified views of boxed regions. Scale bars: 10 μm (main images) and 2 μm (insets). Images are representative of three independent experiments. (B) Quantification of phagosomal Stx2 fluorescence intensity at indicated time points (control: n = 61; 2 h: n = 82; 6 h: n = 56; 12 h: n = 109; 24 h: n = 111). Data represent mean ± s.e.m. from three independent experiments; n.s., not significant. (C) Representative immunoblot showing Stx2 and actin levels in whole-cell lysates from control, zymosan-treated, and *L. major*–infected macrophages. (D) Densitometric analysis of Stx2 levels normalized to actin. Data are mean ± s.e.m. from three independent experiments; n.s., not significant; *P < 0.05. (E) Representative DIC and confocal images of macrophages fed with zymosan particles or infected with wild-type (LmCas9) or GP63-deficient (LmGP63⁻^/^⁻) *L. major* for 6 h, stained for Stx2 (green) and Lamp1 (red). Nuclei were stained with DAPI (blue). Parasite nuclei are indicated by white arrows and zymosan by white asterisks. Insets show magnified regions. Scale bars: 10 μm (main images) and 5 μm (insets). Images are representative of three independent experiments. (F) Quantification of phagosomal Stx2 fluorescence intensity under indicated conditions. Data represent mean ± s.e.m. from three independent experiments; ****P ≤ 0.0001.

To define how *L. major* depletes Stx2, we examined the role of GP63, a GPI-anchored metalloprotease previously shown to target host SNAREs such as VAMP8 (Matheoud et al., 2013). Using CRISPR/Cas9, we generated Cas9-expressing control (LmCas9) and GP63-deficient (LmGP63⁻^/^⁻) *L. major* strains. (Samanta et al., 2025a). Macrophages infected with LmCas9 exhibited marked loss of Stx2 from parasite-containing phagosomes. In contrast, LmGP63⁻^/^⁻ phagosomes retained Stx2 at levels comparable to zymosan-containing phagosomes. (Fig. 2E,F). These data demonstrate that Stx2 depletion is GP63-dependent. Phagosomal Stx2 levels are restored by 24 h p.i., coinciding with promastigote-to-amastigote differentiation, a stage when GP63 expression is strongly downregulated (Hsiao et al., 2008). Amastigotes also remodel the PV by actively acquiring endosomal compartments, which may reintroduce Stx2-positive vesicles (Young and Kima, 2019). Thus, transient Stx2 depletion likely reflects promastigote-specific GP63 activity prior to PV maturation. Mechanistically, we detected no Stx2 cleavage products, suggesting GP63 does not degrade Stx2 directly. Instead, GP63 may promote Stx2 exclusion via protease-independent mechanisms (Brittingham et al., 1999; Lieke et al., 2008; Samanta et al., 2025b). Metalloproteases, including GP63, are known to interact with and influence receptor or signaling protein activity through non-proteolytic mechanisms (Brittingham et al., 1999; Gonzalo et al., 2010). However, further work requires clarifying non-proteolytic phagosomal exclusion of Stx2 by GP63.

### GP63-dependent Stx2 depletion impairs *L. major*–containing phagosome maturation by reduced acquisition of lysosomes and v-ATPase

We next assessed how *L. major* mediated Stx2 depletion affects phagosome maturation. A defining feature of phagolysosome biogenesis is the acquisition of lysosomal components, which is essential for effective microbial killing (Aderem and Underhill, 1999). Consistent with elevated presence of Stx2, phagosomes containing opsonized zymosan robustly acquired the lysosomal marker Lamp1 (Fig. 3A,B). In contrast, phagosomes harboring LmCas9 *L. major* showed significantly reduced recruitment lysosomal marker protein Lamp1, indicative of impaired lysosome fusion. This defect was rescued in phagosomes containing LmGP63⁻^/^⁻ parasites, which acquired Lamp1 at levels comparable to zymosan controls (Fig. 3A). Quantification of Lamp1 fluorescence intensity confirmed a ˵34% reduction in lysosome acquisition by LmCas9 phagosomes, whereas LmGP63⁻^/^⁻ phagosomes were indistinguishable from controls (Fig. 3A). These data indicate that GP63-dependent depletion of Stx2 disrupts lysosomal delivery to *L. major*-containing phagosomes.

**Figure 3.**
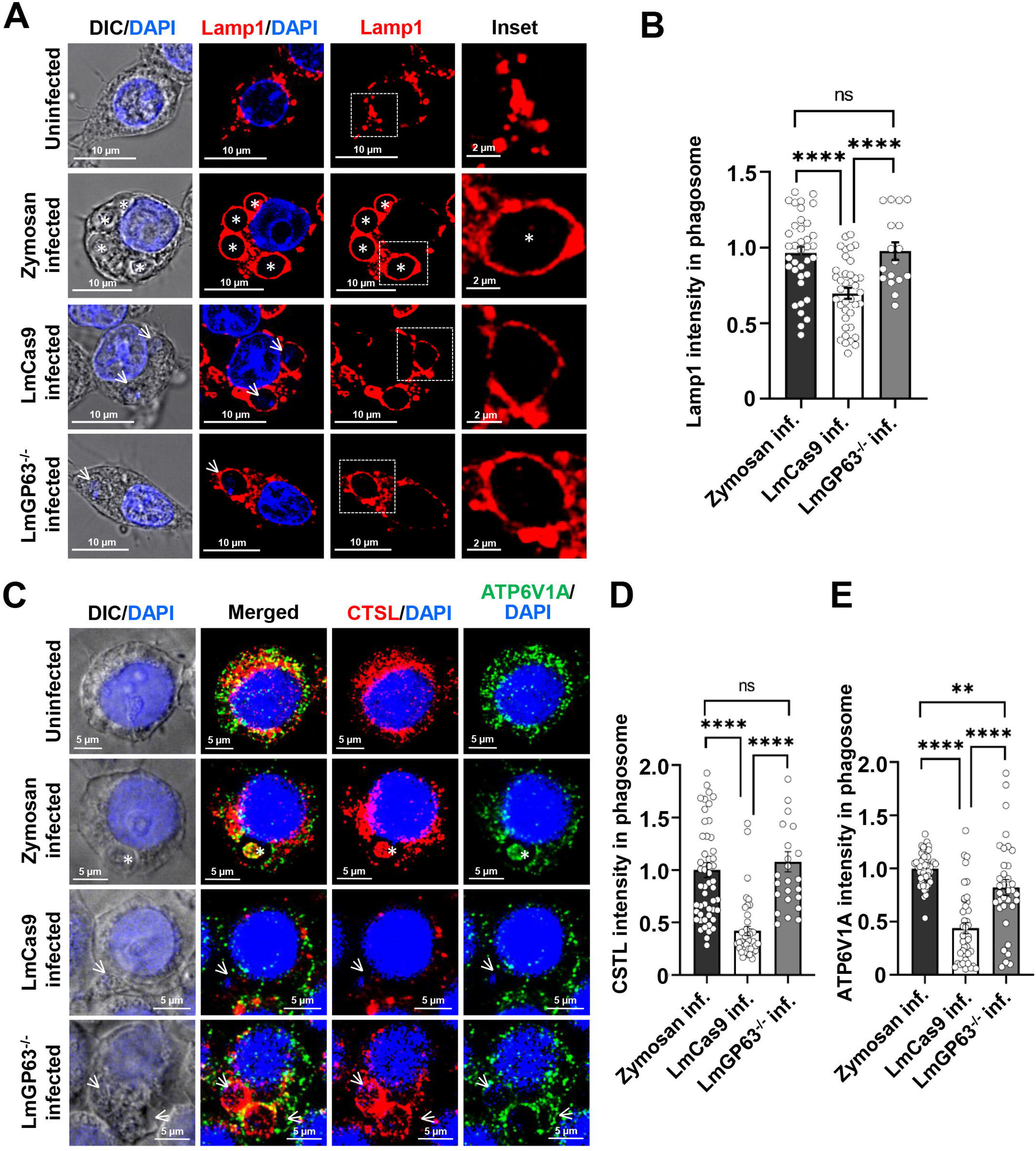
GP63-dependent Stx2 depletion impairs lysosomal and v-ATPase acquisition by *Leishmania major*–containing phagosomes. (A) Representative confocal immunofluorescence images of macrophages under controlled conditions, or following uptake of opsonized zymosan (6 h), or infection with wild-type (LmCas9) or GP63-deficient (LmGP63⁻^/^⁻) *L. major* (6 h), stained for the lysosomal marker Lamp1 (red). Nuclei were counterstained with DAPI (blue). Zymosan particles are indicated by white asterisks. Insets show magnified views of boxed regions. Scale bars: 10 μm (main images) and 2 μm (insets). Images are representative of three independent experiments. (B) Quantification of phagosomal Lamp1 fluorescence intensity under indicated conditions. Data represent mean ± s.e.m. from three independent experiments; n.s., not significant; ****P ≤ 0.0001. (C) Representative confocal images of macrophages under the same conditions as in (A), stained for cathepsin L (CTSL; red) and the v-ATPase subunit ATP6V1A (green). Nuclei were stained with DAPI (blue). Parasite nuclei are indicated by white arrows and zymosan particles by white asterisks. Insets show magnified regions. Scale bars: 10 μm (main images) and 5 μm (insets). Images are representative of three independent experiments. (D–E) Quantification of phagosomal fluorescence intensities for (D) CTSL and (E) ATP6V1A. Data represent mean ± s.e.m. from three independent experiments; n.s., not significant; **P ≤ 0.01; ****P ≤ 0.0001.

To further define the maturation defect, we examined recruitment of cathepsin L [CTSL], a lysosomal protease, and ATP6V1A, a v-ATPase subunit required for phagosomal acidification (Vieira et al., 2002). Compared with control phagosomes, LmCas9-containing phagosomes exhibited markedly reduced recruitment of both CTSL and ATP6V1A (Fig. 3 C–E). This impairment was reversed in LmGP63⁻/⁻ infections, where both markers were efficiently recruited (Fig. 3 C–E). Quantitative analysis revealed ˵57% and ˵60% reductions in CTSL and ATP6V1A fluorescence intensity, respectively, in LmCas9 phagosomes, but no significant difference between LmGP63⁻^/^⁻ and control phagosomes (Fig. 3C-E).

Collectively, these findings demonstrate that GP63-mediated depletion of Stx2 compromises both lysosomal fusion and phagosomal acidification. As a Q-SNARE, Stx2 is central to vesicle docking and fusion with lysosomal compartments (Jahn and Scheller, 2006; Sudhof and Rothman, 2009). Its loss therefore provides a molecular basis for the observed failure to deliver both hydrolytic enzymes and proton pumps to *L. major*-containing phagosomes. While *Leishmania* is known to delay phagosome maturation through lipophosphoglycan and other surface glycoconjugates (Moradin and Descoteaux, 2012), our data implicate GP63-dependent targeting of phagosomal syntaxin as an additional, direct strategy to generate a non-degradative niche. This dual defect in fusion and acidification likely creates an intracellular environment that is permissive for parasite survival but insufficient for antigen processing, with potential implications for immune evasion. Future work will need to address whether Stx2 depletion is specific to *L. major* or represents a conserved virulence mechanism across *Leishmania* species, and whether restoring Stx2 function is sufficient to override GP63-mediated subversion.

### GP63 depletes Stx2 in *L. major* infected BALB/c mice footpad tissues

To determine whether GP63-dependent targeting of Stx2 occurs in vivo, we examined Stx2 levels in footpad tissues from BALB/c mice infected with *L. major* as shown in the scheme (Fig. 4A). Immunofluorescence of cryosections showed robust Stx2 staining in uninfected tissue (Fig 4B). Co-labeling with FcγR, a macrophage marker, confirmed abundant macrophages in the footpad tissues (Fig. S2). At 6 weeks post-infection, LmCas9-infected lesions displayed a marked loss of Stx2 signal within parasite-rich regions (Fig. 4 B). In contrast, LmGP63⁻^/^⁻-infected tissues retained Stx2 levels comparable to uninfected controls (Fig. 4B,C). Quantitative analysis confirmed a significant decrease in Stx2 intensity in LmCas9 infections, which was largely restored in the absence of GP63 (Fig. 4BC). These data demonstrate that GP63 is necessary for Stx2 depletion in vivo.

**Figure 4.**
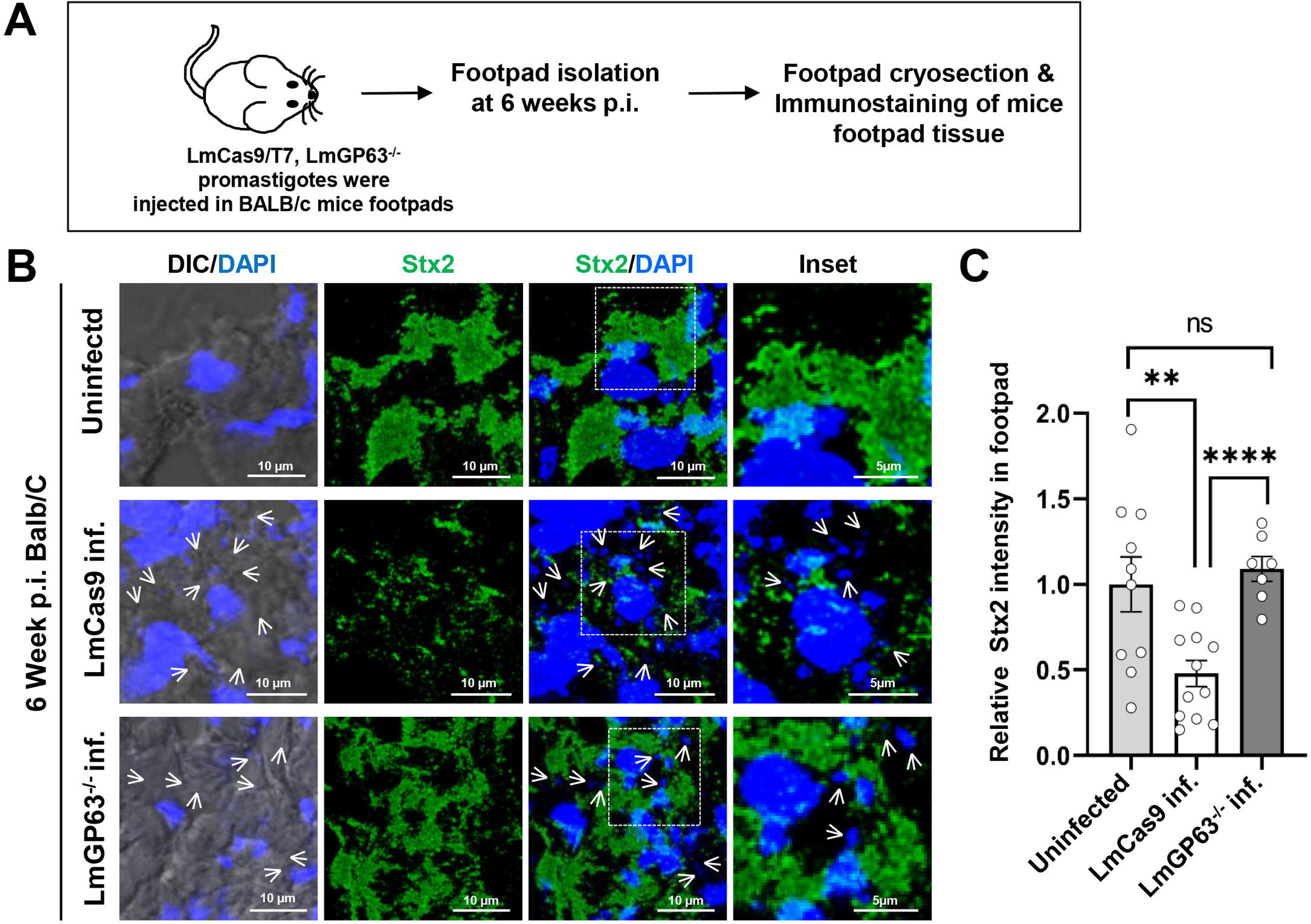
GP63 mediates Stx2 depletion *in vivo* during *Leishmania major* infection. (A) Schematic representation of the experimental design for infection of BALB/c mice with wild-type (LmCas9) or GP63-deficient (LmGP63⁻/⁻) *L. major* parasites. (B) Representative immunofluorescence images of Stx2 (green) in footpad cryosections from uninfected mice or mice infected with LmCas9 or LmGP63⁻/⁻ parasites at 6 weeks post-infection. Nuclei are stained with DAPI (blue), and parasite nuclei are indicated by white arrows. Insets show magnified views of boxed regions. Scale bars: 10 μm (main images) and 5 μm (insets). Images are representative of N = 5 mice per group. (C) Quantification of Stx2 fluorescence intensity in footpad sections. Data represent mean ± s.e.m. (N = 5 mice); n.s., not significant; **P ≤ 0.01; ****P < 0.0001 (unpaired two-tailed Student’s t-test).

These results extend our in vitro findings (Samanta et al., 2025c) to a physiologically relevant infection model and establish the GP63–Stx2 axis as an operative virulence mechanism in vivo. Notably, Stx2 depletion in infected footpad tissue was global and pronounced (Fig. 4B,C), whereas short-term in vitro infection caused only phagosome-specific loss of Stx2 with no change in total cellular levels (Fig. 2). This distinction likely reflects the kinetics of chronic infection. Although amastigotes express lower GP63 levels than promastigotes (Hsiao et al., 2008), sustained exposure over 6 weeks may be sufficient to progressively degrade the cellular Stx2 pool.

Given Stx2’s dual role in regulating phagocytic receptor trafficking and phagolysosome biogenesis (Samanta et al., 2025c), its depletion in situ has several predicted consequences. First, increased surface receptor availability would facilitate uptake of free amastigotes during lesion spread. Second, enhanced transferrin receptor recycling and erythrophagocytosis could improve iron and heme acquisition, both critical for Leishmania growth. Third, loss of Stx2 in bystander macrophages would broadly impair SNARE-dependent delivery of lysosomal hydrolases and v-ATPase, compromising microbicidal capacity at the infection site. Thus, GP63-mediated Stx2 depletion likely remodels the tissue microenvironment to favor parasite persistence. Future work using macrophage-specific Stx2 knockout mice will be needed to dissect the relative contributions of enhanced entry versus defective killing in vivo.

## CONCLUSION

This study identifies Stx2 as a key host regulator of phagosome maturation that is selectively targeted by *Leishmania major* to promote infection. The parasite metalloprotease GP63 depletes Stx2 from pathogen-containing phagosomes, disrupting SNARE-mediated fusion, lysosome and v-ATPase recruitment, and phagolysosome acidification. Stx2 loss enhances parasite uptake and intracellular survival. The GP63-Stx2 pathway operates both in vitro and in vivo, revealing a novel mechanism by which *L. major* subverts host vesicular trafficking to establish a permissive intracellular niche. These findings establish Stx2 as a critical component of antimicrobial defense and a potential target for host-directed therapy in leishmaniasis.

## MATERIALS AND METHODS

### Animals and Macrophages

BALB/c mice (6-8 weeks old) were obtained from the National Institute of Nutrition, Hyderabad. Animals were maintained under specific pathogen-free conditions at the animal facility of IISER Kolkata in accordance with CPCSEA guidelines. All experimental procedures were approved by the Institutional Animal Ethics Committee (IAEC) and conducted in compliance with institutional and national regulations.

Authentic murine RAW 264.7 macrophages (ATCC, catalog no. TIB-71) were maintained in a humidified incubator at 37°C with 5% CO₂. Cells were cultured in Dulbecco’s Modified Eagle’s Medium (DMEM; Thermo Fisher Scientific, catalog no. 12100061) supplemented with 10% heat-inactivated fetal bovine serum (FBS; Avantor, catalog no. 97068-085), 100 U/mL penicillin–streptomycin, and 2 mM L-glutamine.

Stable Control and syntaxin-2 knockdown (Stx2-KD) macrophage cell lines were generated and reported by our group (Samanta et al., 2025c). Briefly, RAW 264.7 macrophages were transduced with lentiviral particles carrying either scrambled control shRNA (Santa Cruz Biotechnology, catalog no. sc-108080) or Stx2-targeting shRNAs (Santa Cruz Biotechnology, catalog no. sc-41327-V). Transduced cells were selected and maintained in complete DMEM supplemented with 5 µg/mL puromycin.

### Parasite Culture

*Leishmania major* promastigotes (strain 5ASKH) were cultured at 26°C in M199 medium supplemented with 23.5 mM HEPES, 10 μg/mL hemin, 150 μg/mL folic acid, 0.2 mM adenine, 120 U/mL penicillin, 60 μg/mL gentamicin, 120 μg/mL streptomycin, and 15% FBS (Avantor, catalog no. 97068-085), as previously described (Dolai et al., 2011). GP63-deficient *L. major* strain (LmGP63−/−) was generated by CRISPR/Cas9 (Samanta et al., 2025a). Cas9- and T7 RNA polymerase-expressing *L. major* cells were maintained under identical conditions with the addition of 50 μg/mL hygromycin. The GP63-deficient (GP63^−/−^) *L. major* strain was cultured in the presence of 20 μg/mL puromycin (Sigma-Aldrich, catalog no. P7255) and 5 μg/mL blasticidin (Thermo Fisher Scientific, catalog no. A1113903) as described earlier (Samanta et al., 2025a).

### Antibodies and Reagents

Primary antibodies used in this study included rabbit polyclonal anti-syntaxin-2 (Stx2; Synaptic Systems, catalog no. 110123, RRID: AB_887849), rabbit monoclonal anti-ATP6V1A (Cell Signaling Technology, catalog no. 39517, RRID: AB_3083783), rabbit anti-Ɣ-actin (Biobharati, catalog no. BB-AB0025), goat polyclonal anti-cathepsin L (CTSL; R&D Systems, catalog no. AF1515, RRID: AB_2087690), mouse monoclonal anti-*Leishmania* GP63 (Thermo Fisher Scientific, catalog no. MA1-81830, RRID: AB_934457), rat monoclonal anti-LAMP1 (CD107a; Thermo Fisher Scientific, catalog no. 14-1071-81, RRID: AB_657532), rat anti-Fcγ (Thermo Fisher Scientific, catalog #14-0161-82, RRID: AB_467133) and human IgG (Sigma-Aldrich, catalog no. I4506, RRID: AB_1163606).

Secondary antibodies included Alexa Fluor 488-conjugated donkey anti-rabbit IgG (Jackson ImmunoResearch, catalog no. 711-545-152, RRID: AB_168465), Alexa Fluor 568-conjugated goat anti-rat IgG (Thermo Fisher Scientific, catalog no. A-11077, RRID: AB_2534121), Alexa Fluor 594-conjugated donkey Anti-Goat IgG (Jackson ImmunoResearch, catalog no. 705-585-003, RRID: AB_156820), and horseradish peroxidase (HRP)-conjugated goat anti-rabbit IgG (Thermo Fisher Scientific, catalog no. 31460, RRID: AB_228341).

Other reagents used in this study includes zymosan A particles (unlabeled, Thermo Fisher Scientific, catalog no. Z2849), RIPA buffer (Santa Cruz Biotechnology, catalog no. sc-24948), latex beads (deep blue-dyed, 0.8 μm; Sigma-Aldrich, catalog no. L1398), puromycin dihydrochloride (Sigma-Aldrich, catalog no. P7255), protease inhibitor cocktail (Sigma-Aldrich, catalog no. P8340), prestained protein ladder (Genetix, catalog no. PG-PMT2922), acetone (Sigma-Aldrich, catalog no. 179124), Dulbecco’s Modified Eagle’s Medium (DMEM; Thermo Fisher Scientific, catalog no. 12100061), fetal bovine serum (FBS; Avantor, catalog no. 97068-085), gelatin (Amresco, catalog no. 9764), glutaraldehyde solution (Sigma-Aldrich, catalog no. G7776), L-glutamine (Thermo Fisher Scientific, catalog no. 25030081), penicillin–streptomycin (Thermo Fisher Scientific, catalog no. 15140122), phosphate-buffered saline (PBS; pH 7.4; Hi-Media, catalog no. TS1101), trypsin-EDTA (0.25%, phenol red; Thermo Fisher Scientific, catalog no. 25200056), paraformaldehyde (Sigma-Aldrich, catalog no. P6148), methanol (Hi-Media, catalog no. MB113), Triton X-100 (Sigma-Aldrich), phenylmethylsulfonyl fluoride (PMSF; Sigma-Aldrich, catalog no. 52332), Laemmli sample buffer (2×; Bio-Rad, catalog no. 1610737), nitrocellulose membranes (0.2 μm; Bio-Rad, catalog no. 1620112), Ponceau S staining solution (Thermo Fisher Scientific, catalog no. A40000279), SuperSignal™ West Pico PLUS chemiluminescent substrate (Thermo Fisher Scientific, catalog no. 34577), skim milk powder (Sigma-Aldrich, catalog no. 70166), and ProLong™ Diamond antifade mountant with DAPI (Thermo Fisher Scientific, catalog no. P36962). All other reagents were obtained from Sigma-Aldrich unless otherwise specified.

### Opsonization of phagocytic particles and phagocytosis

Unlabeled zymosan particles (Thermo Fisher Scientific, catalog no. Z2849) and latex beads (0.8 μm diameter; Sigma-Aldrich, catalog no. L1398) were opsonized with human IgG (Sigma-Aldrich, catalog no. I4506) as described previously (Samanta et al., 2025c). Briefly, zymosan particles were suspended in PBS, briefly sonicated to disperse aggregates, and adjusted to a final concentration of 1 × 10⁸ particles/mL. Latex beads were washed three times with PBS by centrifugation (3,000 × g, 5 min) and resuspended at 5 × 10⁸ particles/mL. Both particles were incubated overnight at 4 °C with 1 mg/mL human IgG for opsonization. Following opsonization, particles were washed three times with ice-cold PBS (3,000 × g, 5 min) and resuspended in complete DMEM for subsequent use.

Phagocytosis assays were performed as previously described (Czibener et al., 2006; Huang et al., 2009; Samanta et al., 2025c). Control or Stx2-depleted (Stx2-KD) RAW 264.7 macrophages (1 × 10^5^ cells per well) were seeded overnight onto 18-mm glass coverslips in 6-well plates (Tarsons, catalog no. 980010). Cells were challenged with opsonized zymosan particles at a ratio of 15:1 (particles:cell) or latex beads at 50:1 (beads:cell; used for phagosome isolation). To synchronize phagocytosis, plates were centrifuged immediately after particle addition (1,000 rpm, 1 min; Hermle Z366K centrifuge with 221.16 rotor) to facilitate uniform particle contact with macrophages, followed by incubation at 37 °C in a humidified 5% CO₂ incubator. Phagocytosis was terminated at indicated time points by replacing the medium with ice-cold PBS. Phagocytic uptake was assessed by fluorescence microscopy, and phagosomes were isolated by density gradient centrifugation as described below.

### Macrophage Infection with *Leishmania major*

RAW 264.7 macrophages were seeded overnight and subsequently stimulated with lipopolysaccharide (LPS; 100 ng/mL) for 6 h. Cells were then infected with *L. major* at a macrophage-to-parasite ratio of 1:30, as described previously (Samanta et al., 2025a). At the indicated time points post-infection, cells were washed with PBS and fixed for subsequent analyses.

### SEM

To assess the initial association of L. major with macrophages, monolayers of control and Stx2-depleted RAW 264.7 cells grown on glass coverslips were infected with parasites at a ratio of 1:30 (cell: parasite) for 10 min. Phagocytic uptake was terminated by the addition of ice-cold PBS. Cells were washed three times with ice-cold PBS to remove unbound and loosely associated parasites and immediately fixed in 2.5% glutaraldehyde (in PBS, pH 7.4) for 1 h at 4 °C. Following fixation, cells were washed twice with PBS and post-fixed in 1% (v/v) aqueous osmium tetroxide for 20 min at room temperature. Samples were subsequently washed with PBS followed by distilled water and dehydrated through a graded ethanol series (30%, 50%, 60%, and 80%) for 5 min each, followed by 100% ethanol for 2 min. Dehydrated samples were dried overnight in a vacuum desiccator and imaged using a Zeiss Supra 55VP scanning electron microscope operating at 5.2 kV.

### Purification of bead and *Leishmania*-containing phagosomes

0.8um OPB containing phagosomes were purified as described previously (Samanta et al., 2025c), adapted from established protocols (Desjardins et al., 1994; Huang et al., 2009; Vinet and Descoteaux, 2009). Briefly, RAW 264.7 macrophages (∼4 × 10⁷ cells per condition) were cultured in 150-mm dishes (Corning, catalog no. CLS430599) and incubated with opsonized latex beads (0.8 μm; 50:1 beads:cell) for 6 h to obtain maturing phagosomes. Cells were washed with warm DMEM to remove unbound beads. Cells were harvested in ice-cold PBS and pelleted (1,200 × g, 4 °C), washed with homogenization buffer (HB; 8.55% sucrose, 3 mM imidazole, 2× protease inhibitor cocktail, pH 7.4), and mechanically disrupted by passage through a 30-gauge needle until >90% lysis was achieved. Post-nuclear supernatants were obtained by centrifugation (1,200 × g, 5 min) and mixed with an equal volume of 62% sucrose. The homogenate was overlaid onto a discontinuous sucrose gradient (62%, 35%, 25%, and 10% in 3 mM imidazole, pH 7.4) and centrifuged (24,100 rpm, 1 h, 4 °C; SW40Ti rotor, Beckman). Phagosomes were recovered from the 10-25% interface, washed with ice-cold PBS, and pelleted at 40,000 × g.

*Leishmania-*containing phagosomes were purified essentially as described previously (Banerjee et al., Cellular Microbiology). Briefly, cells were harvested and homogenized in ice-cold PBS containing protease inhibitors by repeated passage through a 26-gauge needle. The homogenate was initially cleared by low-speed centrifugation at 1,200 × g for 5 min to remove unbroken cells and large debris. The resulting supernatant was further centrifuged at 13,400 x g for 6 min, and the obtained fraction was carefully layered onto a pre-chilled 4-10% sucrose gradient prepared in lysis buffer. Following ultracentrifugation at 30,000 rpm for 45 min, phagosome-enriched fractions were collected from the 4% sucrose layer.

### Western blotting

RAW 264.7 cells or purified phagosomes were lysed in ice-cold RIPA buffer (Santa Cruz Biotechnology, catalog no. sc-24948) supplemented with protease inhibitor cocktail (1×; Sigma-Aldrich, catalog no. P8340). Lysates were sonicated and protein concentrations were determined using a modified Lowry method. Equal amounts of protein were mixed with Laemmli buffer, boiled, resolved by 10-12% SDS-PAGE, and transferred onto nitrocellulose membranes using a semi-dry transfer system (Bio-Rad). Membranes were blocked with 5% skim milk in TBST (TBS containing 0.05% Tween-20) for 1 h at room temperature and incubated overnight at 4 °C with primary antibodies diluted in TBST. After washing (3 × 10 min, TBST), membranes were incubated with HRP-conjugated secondary antibodies (1:4000) for 1.5 h at room temperature. Blots were developed using enhanced chemiluminescence reagents and imaged using a ChemiDoc system (Bio-Rad). For phagosome samples, membranes were stained with Ponceau S following transfer to verify equal loading prior to blocking. Densitometric analyses were performed using ImageJ (v1.54i, NIH). Band intensities were normalized to loading controls, and relative values were calculated against control samples.

### BALB/c mice Infection with *L. major*

As described previously (Samanta et al., 2025a), female BALB/c mice (6-8 weeks old; n = 5 per group) were subcutaneously injected in the left hind footpad with 5 × 10⁶ late stationary-phase *L. major* promastigotes. Control mice received PBS.

### Cryosectioning and immunostaining of mice footpad tissue

At 6 weeks post-infection, mice were euthanized, and infected footpads were harvested and immediately embedded in optimal cutting temperature (OCT) compound. Tissues were frozen at -20 °C overnight and sectioned at 5 μm thickness using a Leica CM1950 cryostat. Sections were collected onto poly-L-lysine–coated glass slides, air-dried, and fixed in 4% paraformaldehyde (PFA) for 10 min, followed by washing with PBS. Sections were permeabilized with 0.2% Triton X-100 in PBS for 15 min, washed twice with PBS, and blocked with 0.2% gelatin in PBS for 30 min. Samples were incubated with primary antibodies (anti-Stx2, 1:100; anti-Fcγ, 1:100) for 1 h at room temperature. After washing (3 × PBS), sections were incubated with appropriate secondary antibodies for 1.5 h at room temperature. Finally, sections were mounted using antifade mounting medium containing DAPI and imaged using a Leica SP8 confocal microscope.

### Immunostaining and confocal microscopy

RAW 264.7 macrophages grown overnight on glass coverslips were washed with PBS and fixed in 4% paraformaldehyde (PFA) for 15 min at room temperature. Following washes with PBS, cells were permeabilized with 0.1% Triton X-100 in PBS for 2 min, washed, and blocked with 0.2% gelatin in PBS for 10 min. Cells were incubated with primary antibodies diluted in blocking buffer for 1 h at room temperature. After washing (3 × PBS), cells were incubated with Alexa Fluor-conjugated secondary antibodies (1:800) for 1 h. Coverslips were washed, mounted using antifade mounting medium containing DAPI (Thermo Fisher Scientific, catalog no. P36962), and imaged. Confocal Z-stack images were acquired at 0.34 μm intervals using a Leica SP8 confocal microscope equipped with a 63×/1.4 NA oil immersion objective. Images were deconvolved using Leica Lightning software.

Fluorescence intensities were quantified using Zen (Carl Zeiss) software and normalized to the mean control signal to obtain relative values.

### Statistical analysis

Statistical analyses were performed using two-tailed unpaired Student’s t-tests in GraphPad Prism (v9.3.0). Data are presented as mean ± s.em. All experiments were independently repeated at least three times. The exact numbers of experiments and the cells or phagosomes quantified are indicated in the respective figure legends.

## ACKNOWLEDGEMENTS

The authors thank the IISER Kolkata core imaging facility (DBT-Builder grant BT/INF/22/SP45383/2022) and acknowledge Mr. Kashinath Sahu for assistance with SEM. We also thank to R.D. lab members for technical support, and Prof. Jayasri Das Sarma for valuable research inputs.

## DATA AND RESOURCE AVAILABILITY

All data reported in this study are available from the lead contact upon request. This study does not report original code. Additional information required for data reanalysis is also available from the lead contact upon request.

## AUTHOR CONTRIBUTIONS

Conceptualization: S.D., S.S.; Data curation: S.S., S.D., R.D; Formal analysis: S.S., S.D., R.D; Funding acquisition: S.D., R.D.; Investigation: S.S., S.D., R.D.; Methodology: S.S., A.P., S.D., R.D.; Project administration: S.D., R.D.; Resources: S.D., R.D.; Software: S.S., A.P., S.D.; Supervision: S.D., R.D.; Validation: S.D., S.S., R.D.; Visualization: S.S., S.D., R.D.; Writing – original draft: S.D., S.S., R.D.; Writing – review & editing: S.S., S.D., A.P., R.D.

## FIGURE LEGENDS

**Figure S1. GP63 reduces total cellular levels of CTSL and v-ATPase components.** (A-B) Quantitative analysis of total fluorescence intensities for CTSL (A) and ATP6V1A (B), derived from the data presented in Figure 3C. Values are expressed as mean ± SEM from three independent experiments. Statistical significance was determined using an unpaired two-tailed Student’s t-test (n.s., not significant; ****p ≤ 0.0001).

**Figure S2. Localization of Stx2 in macrophages within uninfected BALB/c mouse footpad tissue.** (A) Immunofluorescence staining for Stx2 (red) and the macrophage-specific marker CD32 (FcR) (green) in the uninfected footpad cryosections of BALB/c mice. Tissues were harvested at 6 weeks post infection (p.i.). Nuclei were stained with DAPI (blue). Images were acquired with Leica SP8 confocal, 63× objective

