## Supplementary Figures for "*Leishmania major* targets macrophage Syntaxin-2 to impair phagolysosome biogenesis and promote intracellular survival"

### Figure S1 (Related to Fig 2)

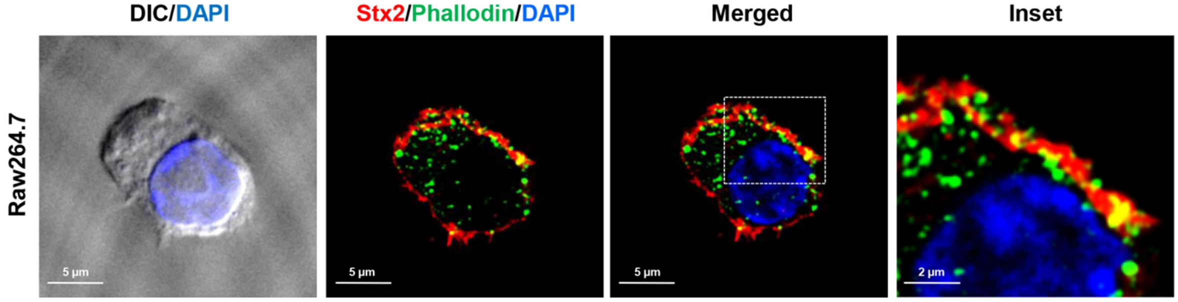


**Figure S1. GP63 reduces total cellular levels of CTSL and v-ATPase components.** (A-B) Quantitative analysis of total fluorescence intensities for CTSL (A) and ATP6V1A (B), derived from the data presented in Figure 3C. Values are expressed as mean ± SEM from three independent experiments. Statistical significance was determined using an unpaired two-tailed Student's t-test (n.s., not significant; ****$p \leq 0.0001$).

**Supplementary File**

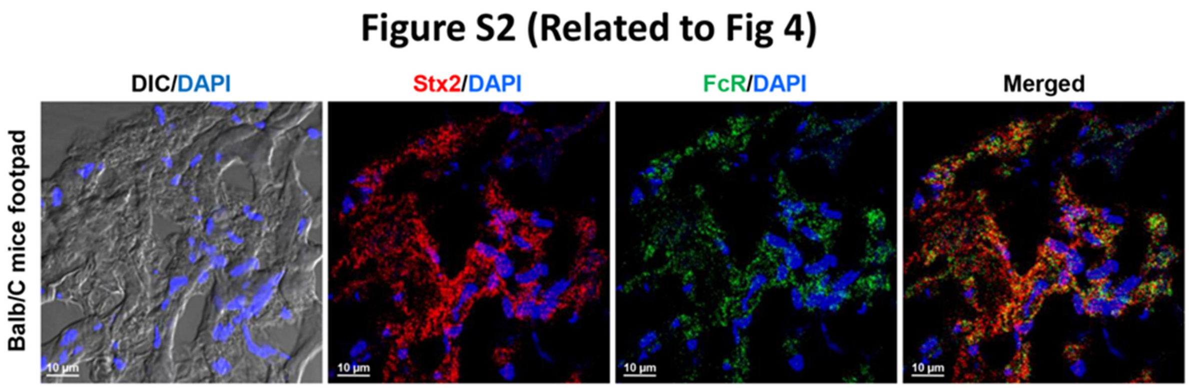


**Figure S2. Localization of Stx2 in macrophages within uninfected BALB/c mouse footpad tissue.** (A) Immunofluorescence staining for Stx2 (red) and the macrophage-specific marker CD32 (FcR) (green) in the uninfected footpad cryosections of BALB/c mice. Tissues were harvested at 6 weeks post infection (p.i.). Nuclei were stained with DAPI (blue). Images were acquired with Leica SP8 confocal, 63× objective
